# AB-free kava enhances resilience against the adverse health effects of tobacco smoke in mice

**DOI:** 10.1101/2024.06.25.599576

**Authors:** Tengfei Bian, Allison Lynch, Kayleigh Ballas, Jessica Mamallapalli, Breanne Freeman, Alexander Scala, Yifan Wang, Hussein Trabouls, Ranjith kumar Chellian, Amy Fagan, Zhixin Tang, Haocheng Ding, Umasankar De, Kristianna M. Fredenburg, Zhiguang Huo, Carolyn J. Baglole, Weizhou Zhang, Leah R. Reznikov, Adriaan W. Bruijnzeel, Chengguo Xing

## Abstract

Tobacco smoke remains a serious global issue, resulting in serious health complications, contributing to the onsets of numerous preventive diseases, and imposing significant financial burdens. Despite regulatory policies and cessation measures aimed at curbing its usage, novel interventions are urgently needed for effective damage reduction. Our preclinical and pilot clinical studies showed that AB-free kava has the potential to reduce tobacco smoke-induced lung cancer risk, mitigate tobacco dependence, and reduce tobacco use. To understand the scope of its benefits in damage reduction and potential limitations, this study evaluated the effects of AB-free kava on a panel of health indicators in mice exposed to 2 – 4 weeks of daily tobacco smoke exposure. Our comprehensive assessments included global transcriptional profiling of the lung and liver tissues, analysis of lung inflammation, evaluation of lung function, exploration of tobacco nicotine withdrawal, and characterization of the causal PKA signaling pathway. As expected, Tobacco smoke exposure perturbed a wide range of biological processes and compromised multiple functions in mice. Remarkably, AB-free kava demonstrated the ability to globally mitigate tobacco smoke-induced deficits at the molecular and functional levels with promising safety profiles, offering a unique promise to mitigate tobacco smoke-related health damages. Further pre-clinical evaluation and clinical translation are warranted to fully harness the potential of AB-free kava in combating tobacco smoke-related harms.

**Graphical Abstract:** 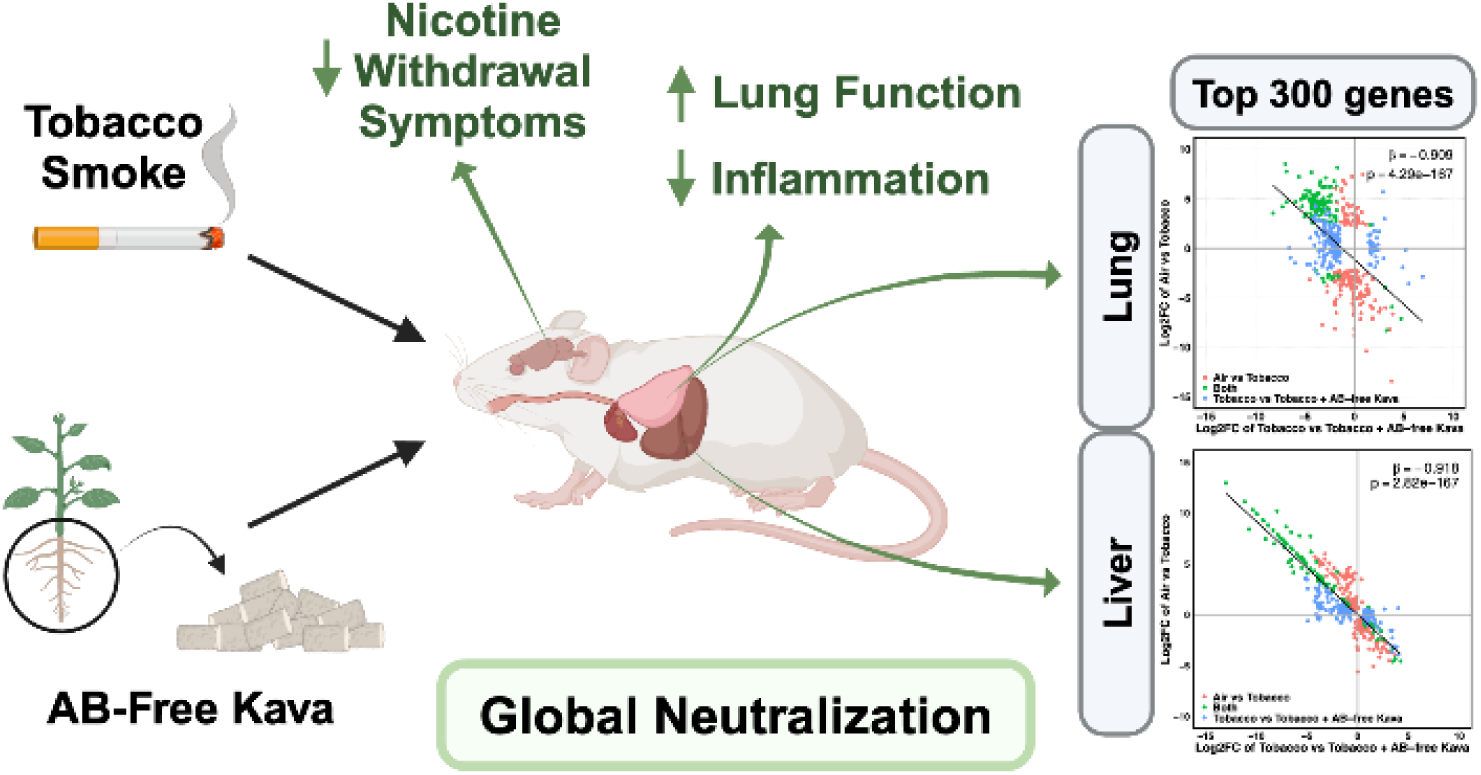

## INTRODUCTION

Despite the development and implementation of various policies and strategies aiming to control tobacco use, tobacco smoking is still a very serious problem in many countries, including the USA, particularly among underserved populations. Chronic smoke exposure induces harm to nearly every organ and is the leading cause of numerous preventable diseases, including lung cancer, cardiovascular diseases, stroke, chronic obstructive pulmonary disease (COPD) and many others. No intervention has been demonstrated to globally counteract the broad harms caused by Tobacco smoke (TS) other than tobacco cessation. Currently, there are about 30 million adult smokers in the USA.^*1, 2*^ Tobacco smoke alone results in half a million premature deaths each year in the USA. Another 16 million American adults live with serious illnesses caused by TS. Indeed, about half of the smokers will die of smoking-related health problems if they fail to quit and tobacco use is estimated to reduce the human life span by 10 – 20 years. At the same time, secondhand smoke exposure is a serious health hazard to many non-smokers, particularly children, causing more than 41,000 deaths per year in the USA. Most smokers are aware of these deleterious health effects and over 50% of them make attempts to quit every year.^*3*^ Tobacco cessation, however, is very challenging, partly due to abstinence-associated withdrawal symptoms, stress, insomnia, and the adverse effects of tobacco cessation therapies, resulting in relapse.^*4–7*^ About 10.0% of American middle and high school students (2.80 million) reported current (i.e., past 30-day) use of tobacco products in 2023,^*8*^ and 22.2% of American middle and high school students (6.21 million) reported ever using any tobacco product.^*9*^ These middle and high school students are at significant risk of continuing tobacco product use in their lifetime. In fact, the percentage of adults who smoke in the USA has persisted at 13 – 15% since 2015^*10*^ and there is little indication that this number will decrease significantly in the near future.^*11*^ Accordingly, smoking-related illness results in approximately $170 billion in direct medical costs and more than $800 billion total lost each year in the USA.^*1*^ Effective interventions, therefore, are urgently needed to ideally help smokers reduce tobacco dependence, achieve cessation, and simultaneously reduce harms imposed by TS exposure. Such interventions are particularly needed for individuals with secondhand and unintentional exposure.

Traditional kava is prepared from the root of *Piper methysticium* as an aqueous suspension and consumed as a beverage among the South Pacific Islanders to reduce stress and improve the quality of sleep.^*12, 13*^ Building on these benefits, organic extract forms of kava had once been used to treat mild and moderate anxiety in Europe.^*14–18*^ A wide range of kava products have also been marketed in the USA as “calming” dietary supplements, which differ substantially in forms, compositions and concentrations^*12*^, and thus have potentially varied benefits and risks. A set of lactones, dominantly detected in kava and thus named kavalactones (Fig. 1, six major ones), are responsible for their relaxing properties with numerous mechanisms proposed.^*19–24*^ Thus, the sum of these six major kavalactones has been used for the standardization of kava products and for the dosage estimation of human consumption.^*12*^ The daily dose of traditional kava is around 750-8,000 mg kavalactones^*13, 25–30*^ while the daily dose for kava clinical use and dietary supplementation is typically 100-300 mg.^*12*^ Kava used during the late 1990s has been associated with rare but severe hepatotoxicity (less than 0.3 case per 10^6^ daily doses), although the exact cause has not been firmly defined. Since traditional kava preparations have been safely used for centuries,^*13, 31–38*^ the World Health Organization (WHO) and others concluded that kava is safe for human consumption with the right materials (the root/stem parts from recommended kava cultivars) and proper preparations.^*13*^ It should be emphasized that traditional kava is a water suspension, whereas kava as a clinical agent was typically an ethanol or acetone extract.^*13*^ Since traditional kava has been safely used for centuries with its daily kavalactone dose 3 – 80 times higher than its clinical dose,^*25–30*^ the reported hepatotoxic risk of kava does not likely derive from the kavalactones and more likely from other ingredients that are enriched in the non-traditional preparations. Flavokavains A and B (Fig. 1) are two chalcone-based compounds more lipophilic than the kavalactones and thus highly enriched in the ethanol or acetone extract forms of kava when compared to the traditional aqueous preparation.^*39*^ They are reported to be the most cytotoxic compounds in kava^*39, 40*^ and induce hepatotoxicity in lab animals alone or in combination with acetaminophen.^*41, 42*^ Compared to the “noble” cultivars typically used for traditional kava preparation, flavokavains A and B are about four times more abundant in low-quality kava cultivars,^*43*^ which are not recommended for human use. During the late 1990s, due to the shortage of noble kava materials and high demand, such low-quality cultivars were used to prepare clinical kava products because of their higher yield and shorter maturation time.^*43*^ Flavokavains A and B, thus, may have contributed to the purported hepatotoxic risk associated with kava. A kava formula, free of flavokavain A and B with a well-defined kavalactone composition mimicking traditional kava preparation, has been developed, named AB-free kava. The six major kavalactones (Fig. 1) account for more than 95% of its mass balance.^*44*^ AB-free kava has demonstrated solid safety profiles in animal models with no signs of adverse risks at dosages equivalent to human 4,000 mg kavalactones per day.^*45–48*^

**Figure 1.**
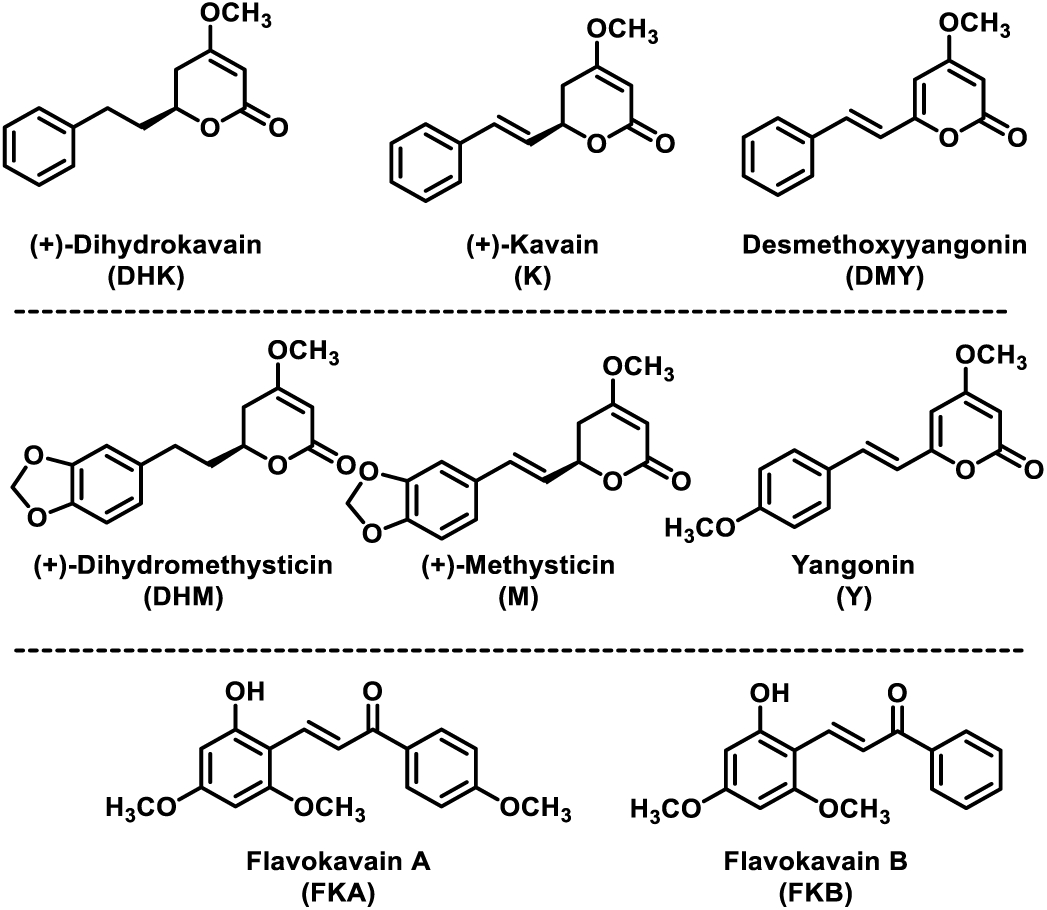
The structures of the six major kavalactones and the two chalcone-based compounds in kava.

Based on an inverse relationship between the cancer incidence rate and the estimated amount of kava consumed among the South Pacific Islanders,^*49–51*^ kava has been hypothesized to reduce human cancer risk. We and others have evaluated and demonstrated that kava could prevent tumorigenesis in multiple pre-clinical animal models.^*46, 52–54*^ For example, kava completely blocked tobacco carcinogen NNK-induced lung tumorigenesis in A/J mice^*52, 53*^ at a dose comparable to the recommended human daily dose of kava as a dietary supplement.^*55*^ Mechanistically, kava reduced NNK-induced DNA damage in the target lung tissues^*52, 53*^ via enhancing carcinogen urinary excretion,^*56*^ both of which can be feasibly and objectively monitored through the quantification of carcinogen metabolites and DNA adducts in the urine. These mechanism-based non-invasive biomarkers greatly enhanced the feasibility of kava’s clinical evaluation. We next performed a pre- and post-AB-free kava one-week AB-free kava supplementation trial among smokers and demonstrated that AB-free kava not only enhanced tobacco carcinogen urinary detoxification but also reduced urinary DNA adducts, substantiating its potential to reduce lung carcinogenesis risk among smokers. In addition, AB-free kava dietary supplementation appeared to reduce tobacco dependence and tobacco use, potentially due to its relaxing properties.^*57*^

Stimulated by these intriguing and beneficial properties of (AB-free) kava, this study investigated the effects of AB-free kava on several distinct biological systems perturbed by TS exposure in pre-clinical mouse models. Unexpectedly, the RNA-Seq results from the lung and liver tissues for the first time revealed that AB-free kava globally neutralized the transcriptional perturbation induced by TS exposure, indicating the unique broad scope of AB-free kava in counteracting various harms induced by TS exposure. This was substantiated by characterizing its effects on an array of different functions known to be perturbed by TS exposure, including lung inflammation, tobacco addiction, and lung function in addition to the potential responsible mechanistic signaling.

## MATERIALS AND METHODS

### Caution

TS is a Class IA human carcinogen and should be handled carefully in well-ventilated fume hoods with proper protective clothing.

### Chemicals and Reagents

The AIN-93M powdered diets were purchased from Harlan Teklad (Cambridgeshire, UK). AB-free kava was prepared by following our reported protocols.^*45*^ Diet supplemented with AB-free kava at the specified dose was prepared by following our reported procedures.^*45*^ 1R6F research cigarettes were purchased from Tobacco and Health Research Institute, University of Kentucky, Lexington, KY. ELISA kits for IL-6 (M6000B) and TNF-α (MTA00B) were purchased from R&D Systems (Minneapolis, MN). BSA standard was purchased from ThermoFisher Scientific (Waltham, MA). The information for antibodies used in this study is included in Table S1. Cotinine and cotinine-d3 standards were purchased from Toronto Research Chemicals (Toronto, Canada). LC-MS grade methanol, water, and formic acid were purchased from Fisher Scientific (Waltham, MA). Oasis MCX cartridges were purchased from Waters (Milford, MA). Acetyl-beta-methacholine-chloride (Sigma Aldrich) was dissolved in 0.9% saline for flexiVent studies. RNeasy Mini Kit (74104) was purchased from QIAGEN (Germantown, MD). 10X TAE Buffer and RiboRuler High Range RNA Ladder were purchased from ThermoFisher Scientific (Waltham, MA) and ethidium bromide was from Sigma-Aldrich (St. Louis, MO).

### Animal study

Mice were purchased from the Jackson Laboratory (Bar Harbor, ME) and maintained in the specific pathogen-free facilities, according to animal welfare protocols approved by Institutional Animal Care and Use Committee at the University of Florida and McGill University. After 7 – 10 days of acclimation, mice were weighed and randomized based on bodyweight into different treatment groups as specified for each round of study detailed below. Mouse bodyweight was measured once a week. For AB-free kava dietary supplementation experiments, daily food intake was estimated twice a week, typically on Monday and Friday, based on the amount of food consumed in each cage, the number of mice housed and the number of days.

### Tobacco smoke (TS) exposure

Tobacco smoke exposure sessions were conducted as described in our previous work.^*58–60*^ Mice were exposed to TS from research cigarettes (1R6F; University of Kentucky, Lexington, NY, USA) via a microprocessor-controlled cigarette smoking machine (model TE-10, Teague Enterprises, Davis, CA) according to the Federal Trade Commission (FTC) protocol with a 2-h exposure daily (9-11 am) from Monday to Friday. Specifically, mice were transferred from their home cage to a clean cage with corncob bedding and a wire top. The cages were placed in the smoke exposure chamber. One smoke exposure system has two large exposure chambers, and each chamber holds up to nine standard mouse cages. The control mice (air-control mice) were placed on a cart in the laboratory during the tobacco smoke exposure sessions. Tobacco smoke was generated with a smoke machine (Teague Enterprises, Davis, CA) and 1R6F research cigarettes (University of Kentucky, Center for Tobacco Reference Products). The cigarettes were smoked using a standardized machine smoking procedure (35 cm^3^ puff volume, 1 puff per min, 2 s per puff, 8 puffs per cigarette). A mixture of mainstream and side-stream smoke was conveyed from the smoke chamber to a mixing and diluting chamber. The smoke was then aged for several minutes and diluted with air to a concentration of ∼100 mg/m^3^ of total suspended particles (TSP). The smoke then passed into the exposure chamber that held the cages with mice and from there into a fume hood to exit the building. The smoke exposure conditions were adjusted by changing the off-time between cigarettes, the number of cigarettes smoked at the same time, and the airflow through the exposure chamber. During each daily smoke exposure session, the TSP count and CO level were measured. The TSP count was determined by gravimetric sampling, and CO levels were determined with a calibrated CO analyzer (Monoxor III, Bacharach, Kensington, PA).

#### Round 1 – 4-week tobacco smoke exposure in A/J female mice

Female A/J mice of six weeks of age were randomized into four groups (n = 6 – 8 mice/group). Mice in Groups 1 and 3 were exposed to filtered air and mice in Groups 2 and 4 were exposed to tobacco smoke, 2 h daily, except Saturday and Sunday, for four weeks. Mice in Groups 1 and 2 were maintained on AIN-M powdered diet while mice in Groups 3 and 4 were maintained on AIN-M powdered diet supplemented with AB-free kava at a dose of 0.75 mg/gram of diet. On Day 4 of Week 2, blood (∼ 40 µL) was collected from mice in Groups 1 and 2 (n = 5) one hour after TS exposure with serum prepared for cotinine quantification. During Week 3, three mice from Groups 1, 2, and 4 were housed in metabolic cages after the 2-h tobacco smoke exposure for 24-h urine collection for tobacco toxicant urinary excretion, which were stored at -80 °C for future analyses. During Week 2 and 4, five mice in Groups 1, 2, and 4 were subjected to somatic withdrawal evaluation as detailed below. All mice were euthanized 22 h after the final tobacco smoke exposure. Four mice each from Groups 1, 2, and 4 were subjected to lung function evaluation as detailed in the Lung Function Measurement section below. For the remaining mice, the blood samples were collected with serum samples processed and stored at -80℃. A portion of the serum samples from mice in Groups 1 and 3 (n = 5 mice/group) were subjected to clinical chemistry analysis (Liver Panel 60405, IDEXX BioAnalytics, Columbia, MO). A portion of the serum samples from Groups 1 – 4 were subjected to TNF-α and IL-6 analyses. Bronchoalveolar lavage (BAL) samples were collected with cells pelleted. The supernatants of the BAL samples were used for TNF-α and IL-6 analysis. The cells from BAL and one lobe of the lung tissues of each mouse were subjected to immune cell profiling, which were detailed in the Immune Cell Profiling section. The rest of the lung and liver tissues were separated into two sections with one section fixed for pathological analysis and one section stored at -80°C. The heart and brain tissues were collected and stored at - 80 °C.

#### Round 2 2-week tobacco smoke exposure in A/J female mice

Female A/J mice of six weeks of age were randomized into three groups (n = 5 mice/group). Mice in Group A were exposed to filtered air and mice in Groups B and C were exposed to tobacco smoke, 2h daily, except Saturday and Sunday, for two weeks. Mice in Groups A and B were maintained on AIN-M powdered diet while mice in Group C were maintained on AIN-M powdered diet supplemented with AB-Free kava at a dose of 0.75 mg/gram of diet. All mice were euthanized 22 h after the final tobacco smoke exposure. BAL was collected successfully for 2, 3, and 4 mice in Groups A, B and C respectively. One lobe of the lung tissues of each mouse and the collected BAL were subjected to immune cell profiling, which was detailed in Immune Cell Profiling section. The rest of the lung and liver tissues were stored at -80°C. The heart and brain tissues were collected and stored at -80°C. The blood was collected with serum prepared.

#### Round 3 – two-weeks tobacco smoke exposure in C57BL/6 male mice

Mice of eight weeks of age were randomized into four groups A-D (n = 5 mice/group) and maintained on standard diet. Mice in Groups 1 and 2 were given 0.2 mL 15% solutol via gavage as control. Mice in Groups 3 and 4 were given AB-free kava at a dose of 80 mg/kg bodyweight via gavage in 0.2 mL 15% solutol two hours before each tobacco smoke exposure. Groups 1 and 3 were exposed to filtered air and Groups 2 and 4 were exposed to TS, 2h daily, except Saturday and Sunday, for two weeks. On the last day, mice were exposed for a single exposure only and were euthanized 24 h after the final tobacco smoke exposure.

### Characterization of AB-free kava in the diet via HPLC

AB-free kava in the diet was extracted and quantified via a published HPLC method with slight modifications.^*61, 62*^ Briefly, AB-free kava diet (200 mg) was placed in a 2-mL Eppendorf tube and mixed with HPLC-grade acetonitrile (1 mL). The sample was vortexed at 3000 rpm for 5 min followed by sonication for 2 min and vortexed again for 2 min. The suspension was centrifuged at 4500 rpm for 30 min. The supernatant was filtered through a 0.22-µm particle filter via a syringe. The diet residue was extracted again through the same process. The filtrates were dried via vacuum centrifugation and reconstituted in HPLC-grade methanol (25 μL) each, which were combined and analyzed via HPLC. An Agilent 100 Series HPLC was used for analysis. The samples (2 µL) were resolved through an Agilent Poroshell 120 SB-C18 column (Agilent, 3.0 x 75 mm, 2.7 µm) maintained at 55°C at a flow rate of 0.40 mL/min with a multi-step gradient previously published using 0.1% formic acid in MilliQ water (mobile phase A) and 7:3 isopropanol: acetonitrile (mobile phase B). Detection λ for the six kavalactones was 240 nm, and for the flavokavains, the λ was 355 nm.

### Cotinine quantification in serum

After thawing on ice, the serum samples in duplicate (2 µL each) were spiked with cotinine-d3 in water (10 pg/µL, 100 pg). Ice-cold methanol (90 µL) was added to precipitate the proteins. The samples were then centrifuged at 4℃ at 13,000 g for 20 minutes. The supernatant was transferred to a new Eppendorf tube and 0.9 mL of 100 mM ammonium acetate (pH = 5) was added. The mixture was then applied to OASIS MCX cartridge (1cc, 30 mg) for solid phase extraction. The cartridges were pre-conditioned with methanol (1 mL) and water (1 mL). The sample was then slowly loaded and washed with 0.5% formic acid in water (2 mL) and methanol (1 mL). The analyte was eluted with 2.5% ammonium hydroxide in methanol (1 mL) and vacuumed to dryness. The analyte was resuspended in 100 mM ammonium acetate in 85% methanol (50 µL) and stored in -20℃ until analysis with 10 µL injected into the LC-MS/MS instrument for analysis. Cotinine from mouse serum was assayed by targeted UPLC-MS2 consisting of a ThermoFisher Vanquish UPLC and a ThermoFisher Altis Plus triple quadrupole. The samples were resolved through a Waters Atlantis HILIC column (150 x 2.1 mm, 3 µm) with a 13-minute gradient with A solvent (0.05% formic acid in 5% acetonitrile/water) and B solvent (0.05% formic acid in acetonitrile) using a 0.2 mL/min flow rate and a linear gradient of 0% B to 80% B followed by a 5-minute equilibration. The mass spectrometer was operated in positive ionization, and detection of the ions was performed in single reaction monitoring (SRM) mode.

### BAL collection

The tracheas of the sacrificed mice were surgically exposed and cannulated with 27-gauge silicon tubing attached to a 23 G needle on a 1-mL syringe. Sterile PBS (1.0 mL) was instilled through the trachea into the lung and withdrawn as the BAL solution. The samples were centrifuged at 300 g for 5 min. The supernatant was collected and stored at -80℃. The cell pellets were used immediately for flow cytometry immune profiling.

### RNA isolation

RNA from mouse lung and liver tissue was isolated using QIAGEN RNeasy Kit following the kit protocol. Briefly, ∼20 mg of tissues from each mouse was homogenized using QIAGEN TissueRuptor homogenizer in Buffer RLT containing 2-Mercaptoethanol (350 µL). The lysate was centrifuged for 15 min at 13,000 g and the supernatant was removed and transferred to a new 1.7 mL microcentrifuge tube. 70% ethanol (350 µL) was added to the lysate and thoroughly mixed by pipetting. Next, the lysate was transferred to a RNeasy spin column and centrifuged for 15 s at 13,000 g; and the flow-through was discarded. The following buffers were subjected to the same sequence in the following order: Buffer RW1 (700 µL), Buffer RPE (500 µL), and Buffer RPE (500 µL). The only difference is the addition of the last Buffer RPE was centrifuged for 2 min instead of 15 s. The RNeasy spin column was placed in a new microcentrifuge tube and the RNA was eluted with RNase-free water (30 µL). The RNA concentration was determined using a BioTek Synergy H1 Microplate Reader with the Take3 microvolume plate. The RNA integrity was checked using a 1% agarose gel run in 1X TAE Buffer stained with ethidium bromide alongside a RiboRuler High Range RNA Ladder.

### RNA-Seq, bioinformatics, and statistical genomics analysis

Paired-end RNA sequencing data was profiled by Novogene Corporation, yielding an average of raw coverage of 25 million pairs of reads. These raw RNA sequencing reads were aligned to the Genome Reference Consortium Mouse Build 38 (GRCm38) using the HiSAT2 software.^*63*^ Duplicate reads identified post-alignment were marked and removed using the Picardtools software. Expression count data were generated using the HTseq software.^*64*^ Lowly expressed genes, defined as having a mean expression level of counts per million reads (cpm) less than 1, were excluded from the lung and liver tissue datasets respectively. Dimension reduction was carried out using Principal Component Analysis (PCA). Differential expression analysis was conducted using a negative binomial model, suited for the count-based nature of RNA-Seq data, as implemented in the R software *edgeR* package.^*65*^ Pathway analysis was conducted using Ingenuity Pathway Analysis (IPA). To further explore the relationship between two lists of differentially expressed genes, we employed the Rank-Rank Hypergeometric Overlap (RRHO) method.^*66*^ The RNA-Seq data has been deposited into Gene Expression Omnibus (GEO) with accession number GSE267842.

### Total protein and albumin quantification in BAL

The concentrations of total protein and albumin in mouse BAL fluid were quantified following the manufacturer’s instruction via the Pierce™ BCA Protein Assay Kit (Thermo Fisher Scientific, MA, USA) and Albumin (BCG) Assay Kit (ab235628, Abcam, MA, USA), respectively.

### Lung pathological evaluation

Mouse lung tissue was formalin-fixed and paraffin embedded for routine staining with hematoxylin and eosin (H&E). H&E stained sections of lung slides from four mice, in each of the four different treatment groups, were evaluated. Under blind conditions, histopathologic changes including karyorrhexis, neutrophilic, lymphocytic, or histiocytic inflammation within large and small airways, alveolated interstitium, and intra-alveolar space were scored on a 0-3 scale (i.e, none, mild, moderate, severe) per anatomic subsite. The totals were combined to generate a single score per study subject.

### TNF-*α* and IL-6 measurements in BAL and serum

TNF-α and IL-6 in BAL fluids and serum samples were measured using ELISA kit (TNF-α: MTA00B; IL-6: M6000B-1) from R&D Systems, according to the recommended protocols from the manufacturer.

### Somatic withdrawal assays

Five mice from Groups 1, 2, and 4 in Round 2 were given mecamylamine (2 mg/kg) via i.p. injection and left to rest for 10 minutes. They were then placed in a clear plastic observation chamber filled with standard bedding and observed for 20 minutes for signs of nicotine withdrawal. Signs included rearing, scratching, digging, forelimb tremors, body shakes, head shakes, abdominal constrictions, jumps, genital licks, and grooming (for more than 3 consecutive seconds). Mice were observed at Week 2 and again at Week 4. All data were analyzed using two-way ANOVA.

### FlexiVent

Pulmonary mechanics were evaluated 20-24 h after the last tobacco smoke administration. Procedures were performed as previously described.^*67*^ Briefly, a tracheotomy was performed in anesthetized mice (ketamine/xylazine/acepromazine) and a cannula (blunted 18 G needle) was inserted into the trachea. Mice were ventilated at 150 breaths/min at a volume of 10 mL/kg of body mass and administered a paralytic (rocuronium bromide). Increasing doses of methacholine from 12.5 to 100 mg/mL^*67*^ were aerosolized using an ultrasonic nebulizer. Anesthetized animals were euthanized at the end of the flexiVent protocol via cervical dislocation.

### Immune cell profiling assays

Mouse tissues were diced into pieces and digested in 10 mL RPMI 1640 containing 1mg/mL collagenase-1A (Sigma-Aldrich) make tissues into single-cell suspensions or thorough homogenates by using semi-automated gentleMACS™ Dissociator and incubate for 45 min at 37 °C on an orbital shaker. After digestion, the samples were filtered through 70 µm cell strainers (Fisher Scientific) to obtain single-cell suspension. Red blood cell lysis buffer (Invitrogen, Carlsbad, CA) was used to lyse residual red blood cells. Bronchoalveolar lavage (BAL) fluid was collected after instilling lungs through the bronchoscope to wash the airways with 1 mL ice-cold PBS. After that cells were wash with PBS and incubated with combination of following antibody; CD11B-PE-dazzle (clone M1/70, 1:200), MHCII I-A/I-E-BB515 (Clone 2G9, BD Biosciences, 1:400), anti-mouse NK1.1-AF700 (clone PK136), anti-mouse CD69-SB436 (clone H1.2F3, eBioscience), antimouse CD3-APC/CY7 (clone 17A2), anti-mouse CD8-BV510 (clone 53-6.7), anti-mouse CD4-BV605 (clone GK1.5), anti-mouse CD279 (clone-PerCP-EF710, eBioscience Inc), anti-mouse CD366-PacBlue (clone B8.2c12), anti-mouse CD62L-BV785 (clone MEL-14), anti-mouse CD45-AF532 (clone 30F.11), anti-mouse CD25-PE-Cy5 (clone PC61), anti-mouse F4/80-BV650 (clone BM8), anti-mouse CD80-BV480 (clone 16-10A1, BD Biosciences), anti-mouse CD11C-PECy7 (clone N418), anti-mouse Ly6G-FITC (clone IA8), anti-mouse Ly6C-BV711 (clone HK1.4), anti-mouse FOXP3-APC (clone FJK16S, 1:50, eBioscience), anti-mouse Granzyme B-BV421 (clone QA18A28FCD, 1:50), antimouse Perforin-PE (clone S16009B), anti-mouse Ki-67-PerCP-Cy5.5 (clone 16A8, 1:50),anti-mouse CD45-AF532 (clone 30F.11), anti-mouse CD170-FITC (clone S17007L), anti-mouse CD68-AF488 (clone FA-11), anti-mouse CD19-PerCP-Cy5.5 (clone 6D5), mouse FcR blocker (anti-mouse CD16/CD32, clone 2.4G2, BD Biosciences) plus LZombie Red™ Fixable Viability Kit (1:500). Flow cytometry was performed by using Cytek Aurora Cytometer (Cytek Biosciences, Fremont, CA) and FlowJo software (BD Biosciences) to analyze data. Most antibodies were used at 1:100 dilution for flow cytometry from Biolegend, unless otherwise specified. Immune cell profiling in Round 3 followed an established procedure.^*68, 69*^

### Western Blotting Assays

20 mg lung tissue was homogenized in 250 µL RIPA buffer and the supernatant was collected after centrifugation at 13,000 g for 15 min at 4 °C. Protein concentrations were determined by the BCA Assay. Lysate (50 µg protein) was electrophoresed on SDS-polyacrylamide gels with proteins transferred onto PVDF membrane. Membranes were blocked with 5% milk in Tris-buffered saline containing Tween-20 before probing with primary antibody according to the manufacturer’s instructions. Subsequently, the membranes were incubated with the corresponding horseradish peroxidase conjugated secondary antibody for 1.5 h. Protein bands were detected by enhanced ECL reagent (Thermo Scientific), and visualized by BioRad Imaging system.

### Statistical Analysis

Data were reported as mean ± standard deviation (SD). One-way ANOVA was used to compare means among multiple treatment groups. For comparison between two groups, two-tailed student *t* test was employed. *p* value ≤ 0.05 was considered statistically significant. All analyses were conducted using GraphPad Prism 4 (GraphPad Software, Inc.).

## RESULTS AND DISCUSSION

### TS exposure characterization

The duration of TS exposure was two to four weeks, as specified, of 2h daily and five days a week for these studies. Total suspended particulate matter (TSP) was 131 ± 35 mg/m^3^ (n = 23) and CO level was 644 ± 111 ppm (n = 23). The level of cotinine in mouse blood 1h after TS exposure was 1.99 ± 0.83 µM (n = 5). These data indicate that the TS regimen used in these studies resulted in TS exposure in mice mimicking the levels and duration among heavy smokers,^*70*^ substantiating the potential physiological relevance of these preclinical studies.

### AB-free kava diet characterization

The composition of AB-free kava and its abundance in the diet was characterized by following a reported HPLC method on the six major kavalactones with slight modifications.^*61, 62*^ The six major kavalactones, the main ingredients in AB-free kava, accounted for more than 95% of the mass balance based on the area estimation of all detectable ingredients at the wavelength of 240 nm, consistent with our previous characterization.^*45*^ The total amount of these six major kavalactones was 725 ± 9 µg/gram of diet for Round 1 and 2, comparable to the initial dose design (750 µg/g of diet). It included methysticin (28.0 ± 0.2 µg/gram of diet), dihydromethysticin (84 ± 1 µg/gram of diet), kavain (285 ± 4 µg/gram of diet), dihydrokavain (251 ± 3 µg/gram of diet), yangonin (24.1 ± 0.2 µg/gram of diet), and desmethoxyyangonin (52.9 ± 0.7 µg/gram of diet) respectively (Fig. S1). Based on the estimated 2-g daily food intake for mice,^*71*^ the dosage of AB-free kava in mice in this study is equivalent to ∼ 450 mg kavalactones daily for a human of 75-kg bodyweight, extrapolated from the body-surface area dose estimation.^*72*^ Such a dose is slightly higher than the recommended dietary supplementary dosages of kava (up to 300 mg kavalactones daily) but well below the estimated daily kavalactone dosages among traditional kava users (750 – 8000 mg kavalactones daily).^*12, 73*^

### Safety measures

The time-course mouse bodyweight changes during the experimental 4-week period in Round 1 showed that tobacco smoke significantly suppressed mouse growth (Fig. S2). This is consistent with previous literature reports about the effects of tobacco smoke on mouse growth.^*74–76*^ Mice supplemented with AB-free kava in the diet showed similar bodyweight changes in comparison to those with control diet (Fig. S2). Similar food intake was observed among mice in different treatment groups (data not shown). Alkaline phosphatase (ALP), alanine transaminase (ALT), aspartate transaminase (AST), total protein, creatinine kinase, and total and free bilirubin in the terminal serum samples were quantified from mice with and without AB-free kava dietary supplementation (n = 5 mice/group). The 4-week AB-free kava dietary supplementation caused no increase in any of these measurements. These data overall support the promising safety profiles of AB-free kava under the current treatment regimen in A/J mice, again consistent with our previous observations.^*45*^

### RNA-Seq profiling of the lung and liver tissues

The RNA-Seq raw data among different groups in the same round of animal study were similar in quality and quantity, supporting their comparative analyses. Principal Component Analysis (PCA) was first performed to evaluate the global effects of different treatment regimens, using all mRNA data obtained from the RNA-Seq analysis. Among the four groups of mice in Round 1 (n = 4 mice/group), no clear group clustering was observed based on the global PCA analysis (Fig. 2A). Such a result indicates that the extent of transcriptional perturbations in the lung tissues introduced by tobacco smoke exposure and/or AB-free kava under the 2-week regimens were generally mild that they did not cause significant global separations among mice with different treatment regimens. This is indeed consistent with the fact that acute and semi-chronic TS exposure has no immediate and obvious detrimental effects on human health and that it in general requires long-term TS exposure to introduce pathological and health changes. Similarly, AB-free kava also did not result in dramatic transcriptional perturbation, which is in alignment with the safe and chronic use of kava in its traditional settings as well. Similar results were observed in the mouse liver tissues in Round 2 (Fig. 2B). We next focused our analyses on those genes significantly perturbed by TS exposure or TS + AB-free kava. We identified 1100 genes in the lung tissues and 999 genes in the liver tissues, that were significantly perturbed by TS in comparison to filtered air (*p* < 0.05). Identified genes were then subjected to IPA analysis to determine the top signaling pathway candidates being perturbed by TS exposure. Similarly, 576 genes in the lung tissues and 1058 genes in the liver tissues were significantly perturbed by TS + AB-free kava in comparison to TS exposure alone. These genes were also used to determine the top signaling pathway candidates via IPA. The relationship between activation z-scores of the top 40 identified pathways (based on statistical significance ranking, not all pathways have a defined z-score) were explored. Strikingly, a strong negative correlation was observed for both lung (correlation coefficient = -0.657, *p* = 6.44e-05, Fig. 2C) and liver tissues (correlation coefficient = -0.737, *p* = 1.22e-08, Fig. 2D). Similar analyses were performed using the top identified 300 genes in both comparisons. They also revealed a strong negative correlation in terms of log2 fold changes for both lung (correlation coefficient = -0.909, *p* = 4.29e-167, Fig. 2E) and liver tissues (correlation coefficient = -0.918, *p* = 2.82e-167, Fig. 2F). These mRNA profiling results suggest that the transcriptional perturbation caused by TS exposure appeared to be globally neutralized upon AB-free kava exposure, for the first time implying that AB-free kava has the potential to holistically address multiple harms caused by TS exposure. Detailed pathway analysis and validation studies are ongoing, which is not within the scope of the current report. Several functions well known to be perturbed by TS exposure, including lung inflammation, nicotine addiction, and lung function, were thus characterized in these studies to assess the global neutralizing potential of AB-free kava on the effects introduced by TS exposure.

**Figure 2.**
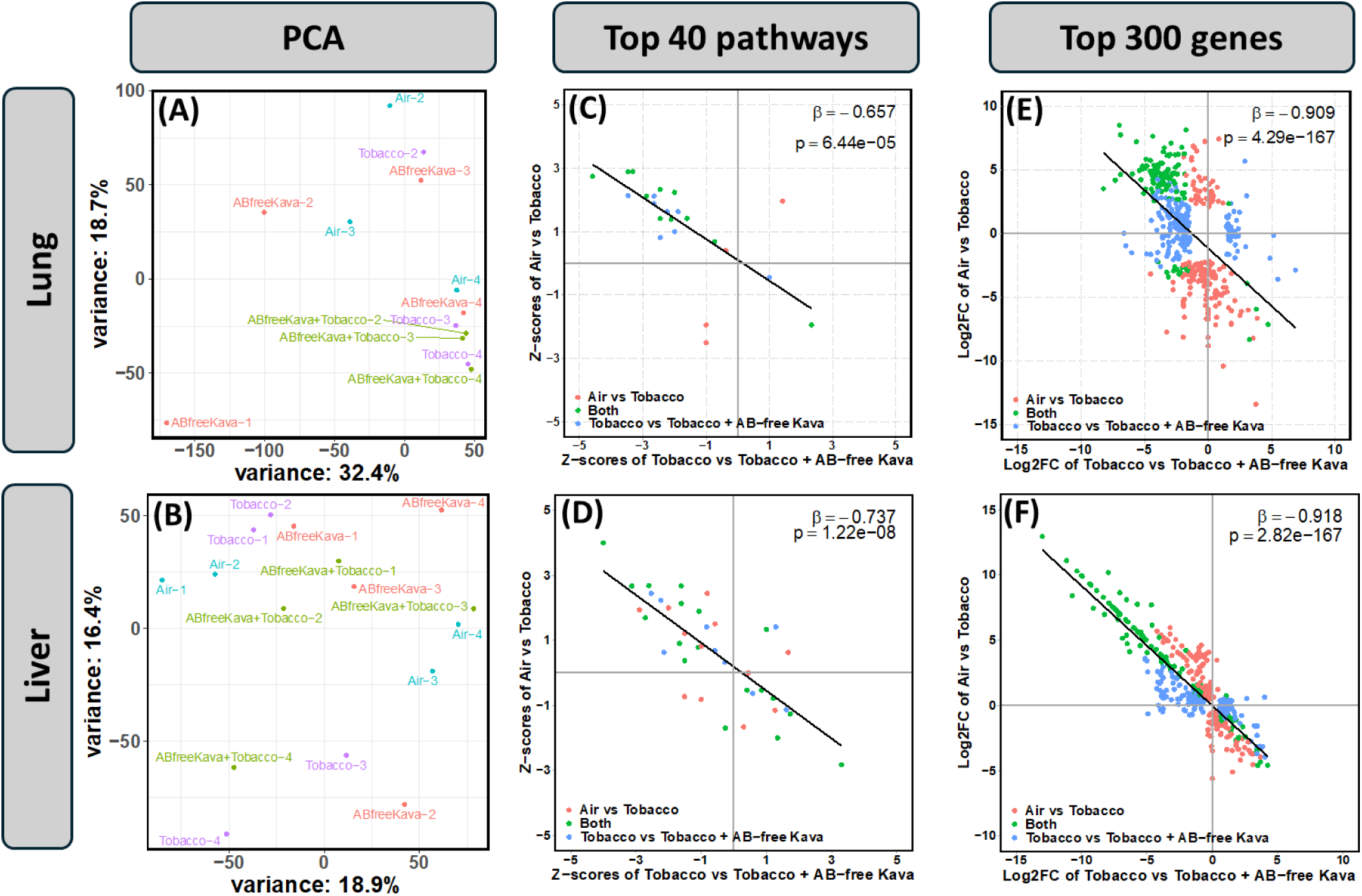
The RNA seq results of the lung and liver tissues from mice with different treatment regimens. Air: Filtered Air; TS: tobacco smoke; and AB-free Kava: AB-free kava. Figure A and B display the principal component analysis (PCA) of the RNASeq data, showing the first two principal components (PCs) for individual mice in lung and liver tissues, respectively. The variance explained by each PC is indicated on the x-axis and y-axis labels. Figure C and D illustrate the correlation of the activation z scores of the top 40 signaling pathways perturbed by TS in comparison to Air (y-axis), and by TS + AB-free kava in comparison to TS (x-axis). β represents the regression coefficient of the scatter plot. These pathways are categorized into three groups: those significant only when comparing TS to Air, those significant only when comparing TS + AB-free kava to TS, and those significant in both comparisons. Figure E and F show the correlation of log2Fold Change among the top 300 genes perturbed by TS in comparison to Air, and by TS + AB-free kava in comparison to TS.

### Lung inflammation

TS is well known to induce lung inflammation, which may have contributed to the development of a wide range of lung diseases, particularly COPD and lung cancer. We thus profiled the immune cells in different tissues – including BAL, lung, spleen, thymus and bone marrows – from TS and/or AB-free kava treated mice after two weeks or four weeks of treatment, using CD11b^+^Ly6C^+^Ly6G^+^ population as neutrophils (Fig. 3A). In both two weeks and four weeks treatment groups, we found that TS significantly increased neutrophil cells in BAL while AB-free kava reversed it (Fig. 3B-3C, BAL-Neu); whereas AB-free kava slightly increased neutrophils located within the lung tissues after excluding BAL fractions (Fig. 3B-3C, Lung-Neu). Blood neutrophils were unchanged (Fig. 3B, Blood-Neu). We did not observe significant changes in other immune cells we have profiled, including populations in the T cells, B cells, and other myeloid cells (data not shown). The consistent reverse distribution pattern of neutrophils within the BAL and lung indicates that TS exposure induces mild lung inflammation such that neutrophils from the lung migrate to the BAL to fight against the inflammation while AB-free kava may reduce the inflammatory activity via reducing the infiltration of neutrophils from the lung into the BAL. No significant changes in immune cell profiles were observed in the spleen, thymus and bone marrow (data not shown), indicating that TS under the current treatment regimen induces inflammation mainly in the lung tissues.

**Figure 3.**
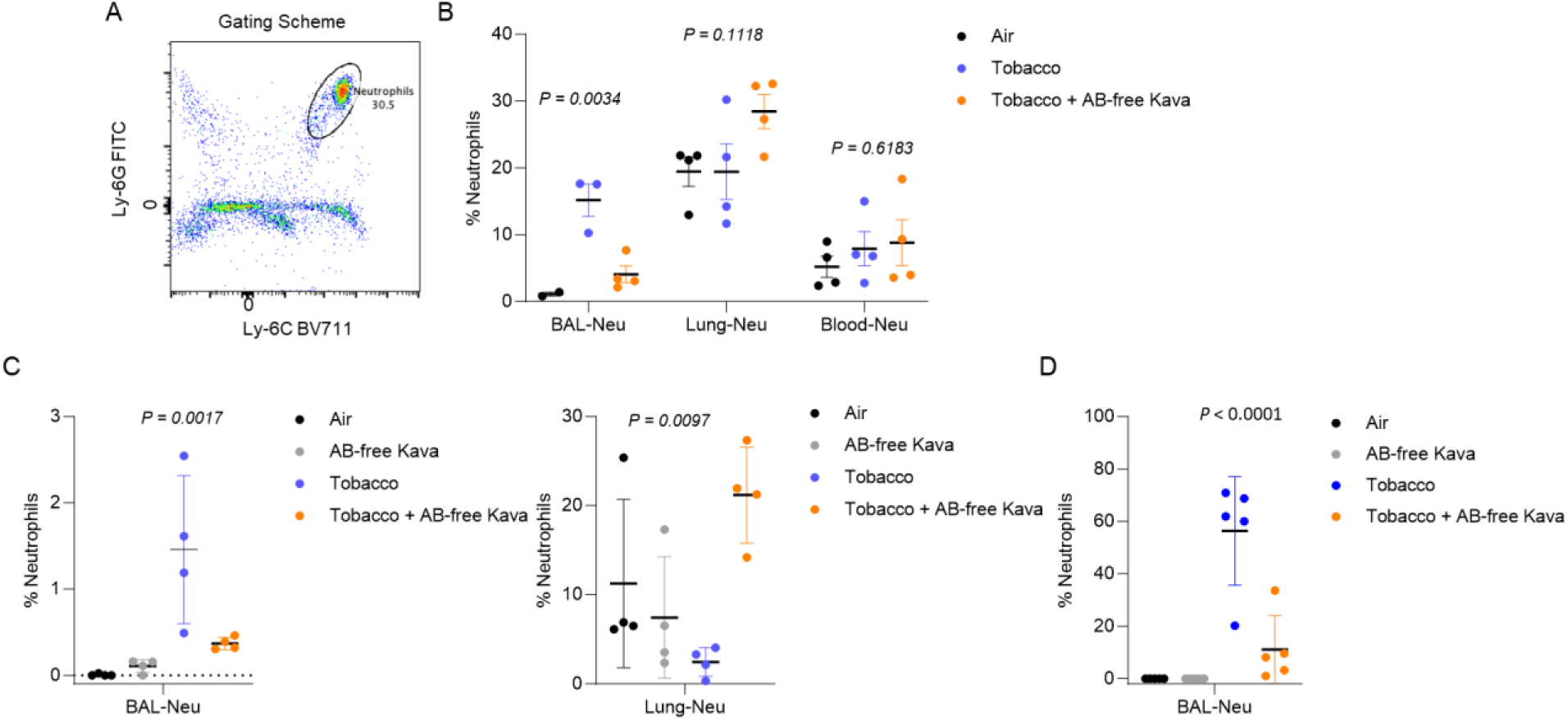
The effect of TS and/or AB-free kava on proinflammatory immune cells in the BAL and lung tissues. **A-C.** Flow cytometry analysis of neutrophils in the BAL and lung tissues. (A) Gating schematic for neutrophils showing Ly6C^+^Ly6G^+^ population. The effect of TS exposure and/or AB-free kava via gavage on neutrophils in mouse BAL, after (B) two weeks or (C) four weeks of TS and/or AB-free kava (n = 4 mice/group). **D.** BAL smear-based pathological determination of neutrophil numbers in BAL after two weeks of TS and/or AB-free kava (n = 5 mice/group).

To confirm the BAL neutrophil number and frequency, we also collected BALs from the 4 groups of mice after two weeks of treatment, and performed Giemsa staining on BAL smears.^*69*^ Two weeks of TS exposure significantly increased neutrophil population in BAL; whereas AB-free kava reduced such an increase by ∼70% (Fig. 3D), validating the anti-inflammatory potential of AB-free kava against TS exposure via reducing BAL neutrophil accumulation.

Consistent with the neutrophil population changes, significant increases in TNF-α (Fig. 4A) and IL-6 (Fig. 4B) were observed in mouse BAL and serum samples upon TS exposure while AB-free kava significantly reduced the levels comparable to those in the control mice. With respect to the epithelial cell integrity of the lung tissues, the total protein concentration (Fig. 4C) and albumin concentration (Fig. 4D) in the BAL remained similar among samples from mice with different treatments. Consistently, no significant pathological lesions were detected in the lung tissues upon current tobacco smoke exposure regimen (Fig. 4E, representative lung tissues for the upper panel: TS group; lower pane: TS + AB-free kava group). These data overall indicate that TS exposure has not significantly compromised the integrity of the lung epithelial cells under the current treatment regimens.

**Figure 4.**
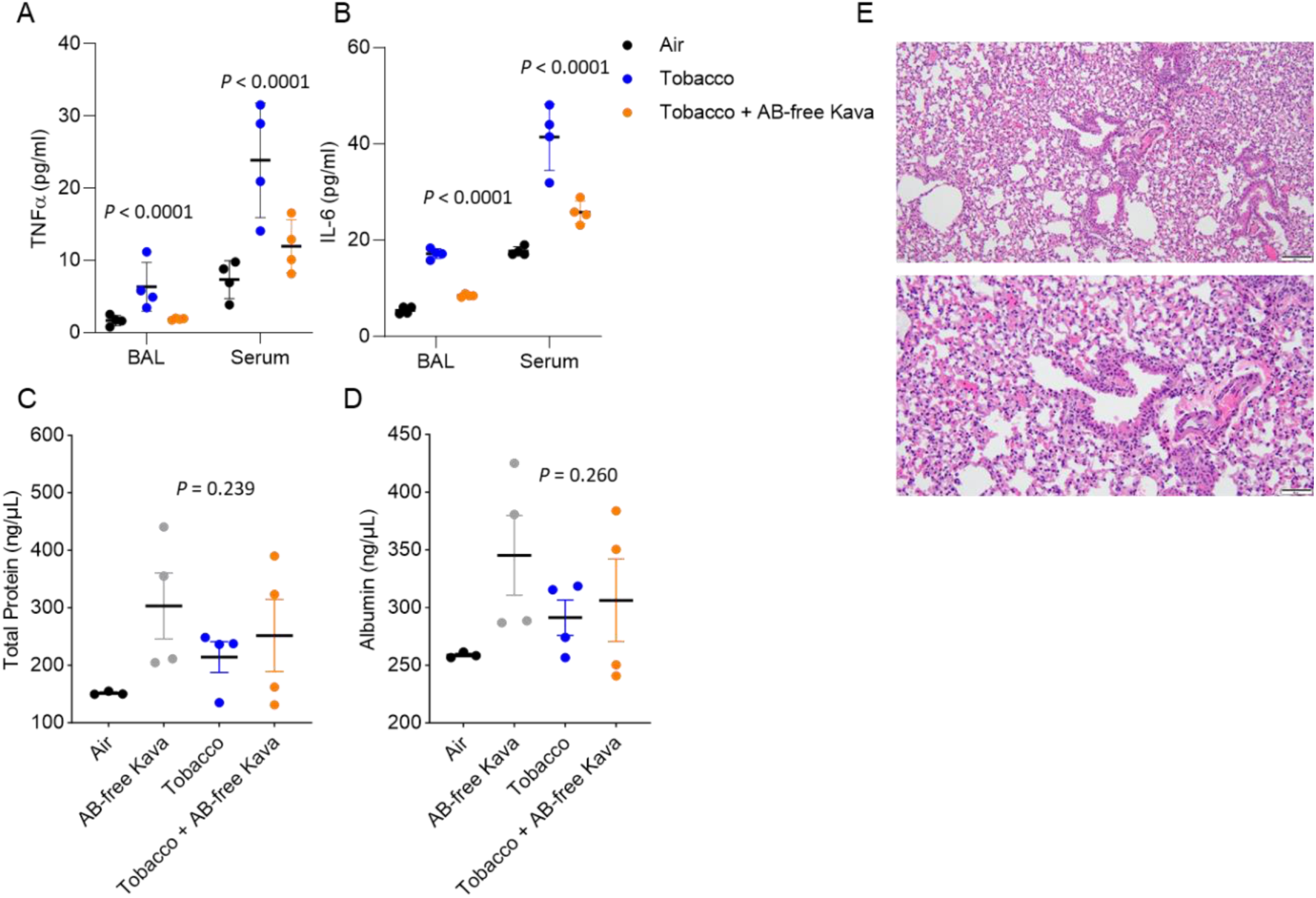
The effect of TS and/or AB-free kava on proinflammatory cytokines and lung tissues. **A-B.** (A) TNF-α or (B) IL-6 in mouse BAL or serum samples of different treatment groups after 2 weeks of TS and/or AB-free kava (n = 4 mice per group). **C-D**. The effects of tobacco smoke and/or AB-free kava via dietary supplementation on (C) total protein and (D) albumin levels in mouse BAL after 2 weeks of TS and/or AB-free kava (n = 4 mice per group). **E.** Representative lung tissue pathological images from mice from different groups with no obvious lesions related to different treatment regimens (top, TS treated lung; bottom, TS + AB-free Kava).

### Somatic withdrawal symptom evaluation

The results of our pilot one-week kava supplementation trial indicate that kava has the potential to reduce tobacco dependence.^*57*^ We thus evaluated the potential of AB-free kava for tobacco cessation in this tobacco smoke model by assessing somatic withdrawal signs. Somatic withdrawal signs were counted after one week and 4 weeks of tobacco smoke exposure (Fig. 5). Mecamylamine precipitated somatic withdrawal signs increased over time (Time F1,12 = 15.521, *p* = 0.002). Treatment with AB-free Kava prevented the mecamylamine-induced increase in somatic withdrawal signs in the smoke exposed mice over time (Time x Treatment F2,12 = 15.395, *p* < 0.001; Treatment F2,12 = 17.943, *p* < 0.001). The post hoc test showed that the tobacco smoke-exposed mice displayed more withdrawal signs than the air-control mice during weeks 1 and 4. The tobacco smoke-exposed mice displayed more withdrawal signs during week 4 than during week 1. Furthermore, the tobacco mice that were treated with AB-free kava did not display more somatic withdrawal signs than the air-control mice during weeks 1 and 4.

**Figure 5.**
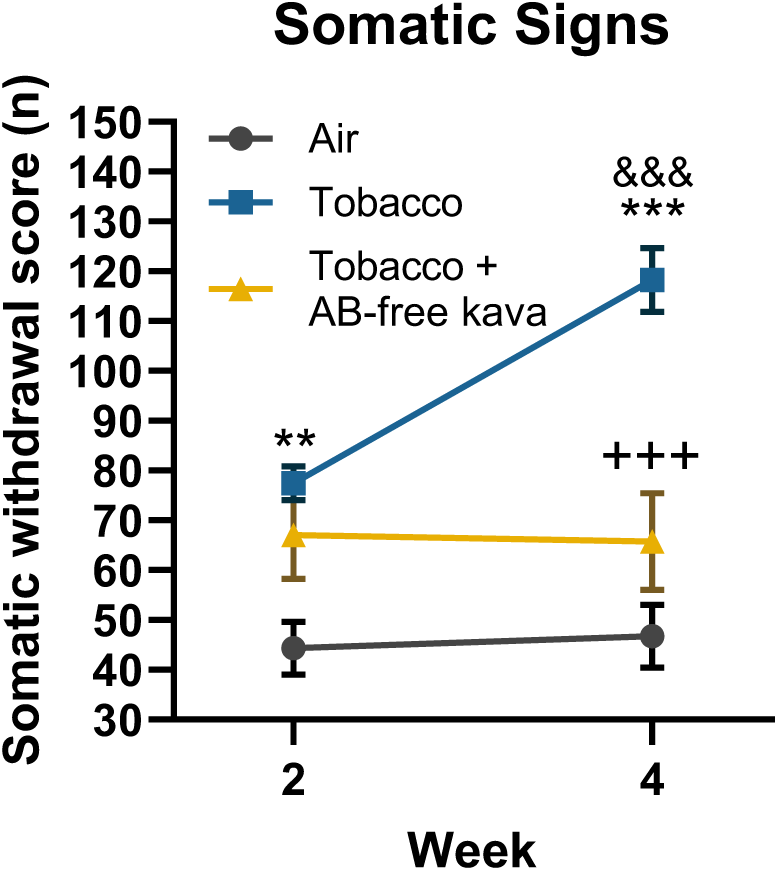
AB-free kava diminishes somatic withdrawal signs in tobacco smoke-exposed mice. Asterisks indicate more signs compared to the air group during the same week. Plus signs indicate fewer signs compared to the tobacco group during the same week. Ampersand signs indicate more signs compared to the tobacco group during week 2. **, *p* < 0.01; ***, +++, &&& *p* < 0.001.

We also determined the effects of AB-free kava on the individual somatic withdrawal signs (Table 1). Grooming did not change over time and was not affected by tobacco smoke exposure or AB-free kava (Time F1,12 = 1.846, NS; Time x Treatment F2,12 = 1.208, NS; Treatment F2,12 = 1.617, NS). Digging increased over time but this was not affected by tobacco smoke exposure or AB-free kava (Time F1,12 = 5.441, *p* = 0.038; Time x Treatment F2,12 = 0.286, NS; Treatment F2,12 = 0.341, NS). Rearing increased over time and treatment with AB-free kava prevented the nicotine withdrawal-induced increase in rearing (Time F1,12 = 6.534, *p* = 0.02; Time x Treatment F2,12 = 16.183, *p* < 0.001; Treatment F2,12 = 25.947, *p* < 0.001). The posthoc test showed that the mice in the tobacco group displayed more rearing during week 4 than during week 1. Furthermore, the mice treated with AB-free kava displayed less rearing than the mice in the tobacco group during week 4.

**Table 1.** Effect of AB-free kava on individual somatic withdrawal signs.

|  |  | Week 2 | Week 4 |
| --- | --- | --- | --- |
| Grooming | Air | 13.8 ± 2.3 | 9.6 ± 1.3 |
|  | Tobacco | 11.4 ± 2.0 | 11.8 ± 1.2 |
|  | Tobacco + AB-free Kava | 9.0 ± 2.1 | 7.8 ± 1.7 |
| Digging | Air | 13.4 ± 3.3 | 17.8 ± 3.1 |
|  | Tobacco | 16.2 ± 3.4 | 22.6 ± 1.9 |
|  | Tobacco + AB-free Kava | 12.0 ± 4.4 | 21.8 ± 7.0 |
| Rearing | Air | 16.8 ± 3.1 | 18.6 ± 4.8 |
| | Tobacco | 49.6 ± 5.2** | 83.2 ± 5.7**, \$\$\$ |
|  | Tobacco + AB-free Kava | 46.0 ± 7.6** | 35.6 ± 5.9<sup>++</sup> |
| Genital licks | Air | 0.4 ± 0.2 | 0.8 ± 0.5 |
|  | Tobacco | 0.2 ± 0.2 | 0.6 ± 0.4 |
|  | Tobacco + AB-free Kava | 0.0 ± 0.0 | 0.6 ± 0.4 |

Asterisks indicate more signs compared to Air group during the same week, plus signs indicate less signs compared to Tobacco group during the same week. Dollar signs indicate more signs compared to Tobacco week 2.

These data overall are consistent with our pilot clinical trial results, supporting the potential of AB-free kava to facilitate tobacco smoking cessation, potentially due to its relaxing and sleep improving properties. We further hypothesize that AB-free kava reduced somatic withdrawal signs by diminishing noradrenergic transmission in the brain. AB-free kava has been shown to inhibit NE-induced intracellular calcium influx in H1299 cells by antagonizing β-adrenergic receptor signaling.^*77*^ Interestingly, in our previous work, we showed that the β-adrenergic receptor antagonist propranolol decreased the total number of somatic signs associated with nicotine withdrawal.^*78*^ These findings suggest that AB-free kava diminishes nicotine withdrawal by diminishing noradrenergic transmission via β-adrenergic receptor blockade. Since abstinence-associated withdrawal, stress, and sleep issues contribute to the limited success rate of tobacco cessation by current medications, AB-free kava may provide a unique and complementary opportunity to help addicted smokers quit.

### Lung functional analyses

Lastly, given the detrimental effects of TS exposure on lung functions, we evaluated the effects of 4-week TS exposure with and without AB-free kava dietary supplementation on the mouse lung function via flexiVent. We first examined the impact of TS administration on airway resistance using single frequency forced oscillation technique. We found no effect of treatment on basal airway resistance (Fig. 6A). When examining normalized airway resistance, we observed that methacholine increased airway resistance in response to increasing concentrations of methacholine in all groups (Fig. 6B). However, the magnitude of this increase was diminished in the TS + AB free-kava group compared to the Air and TS groups (Fig. 6B; Treatment, F2, 40 = 3.661; *p* = 0.035). Performing a post hoc analysis corrected by Sidak’s multiple comparison test revealed no significant differences between groups at any dose of methacholine.

**Figure 6.**
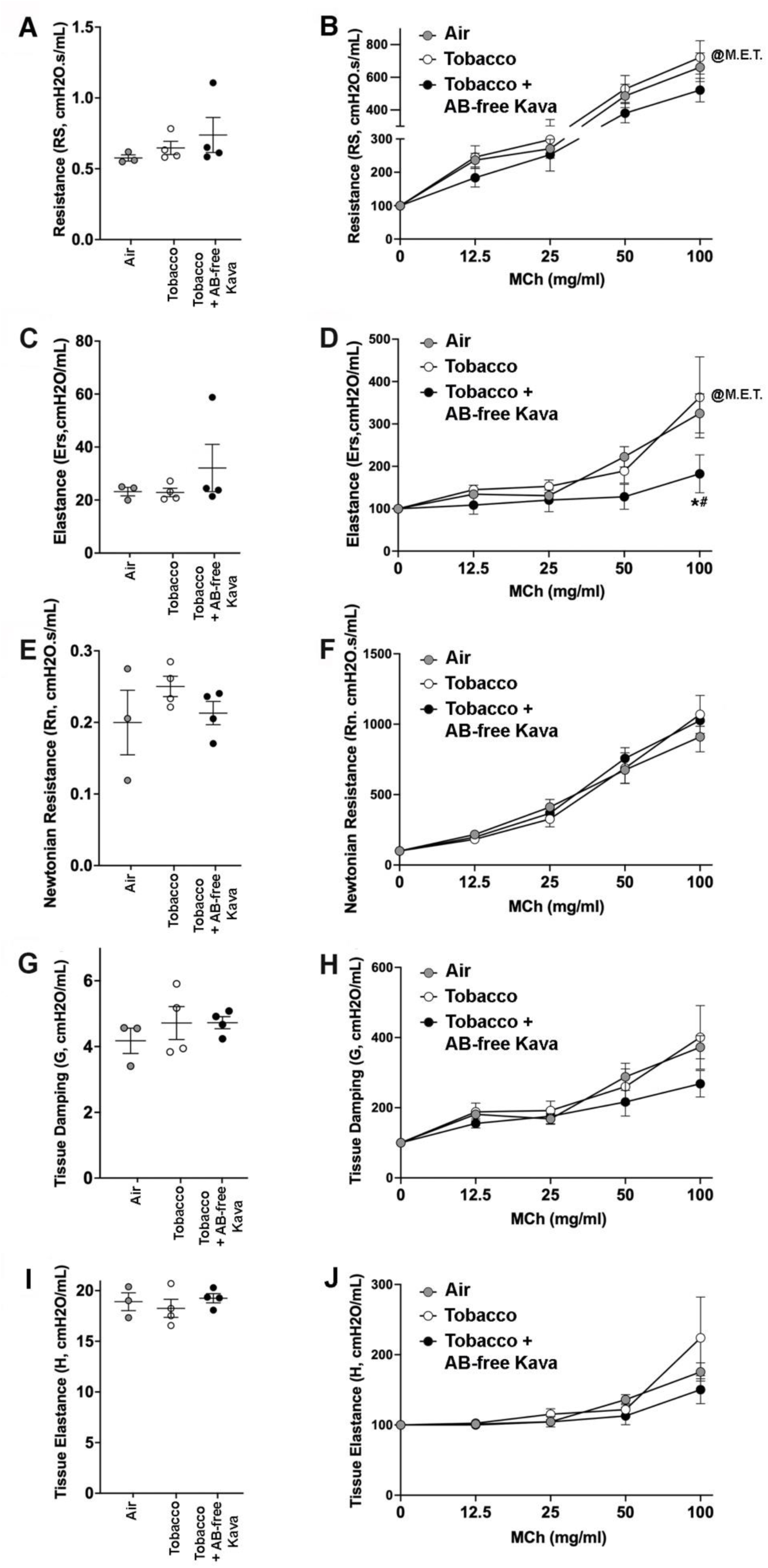
The effects of tobacco smoke exposure with/without AB-free kava dietary supplementation on lung function characterization. **(A)** Basal airway resistance. (**B**) Normalized resistance in response to methacholine in male airways. (**C**) Basal airway elastance. (**D**) Normalized elastance in male airways in response to methacholine. * = compared Air. #; compared to TS. (**E**) Basal airway Newtonian resistance. (**F**) Normalized Newtonian resistance in male airways in response to methacholine. (**G**) Basal tissue damping. (**H**) Normalized tissue damping in male airways in response to methacholine. (**I**) Basal tissue elastance. (**J**) Normalized tissue elastance in male airways in response to methacholine. Air (n = 3), Tobacco (n = 4); Tobacco + AB-free kava (n = 4). Abbreviations: M.E.T., main effect of treatment.

Single frequency forced oscillation also provides airway elastance measurements. We found no effect of treatment on basal elastance values (Fig. 6C). However, treatment modified the normalized airway elastance response to methacholine (Fig. 6D; Treatment, F2, 40 = 4.674; *p* = 0.015). Post hoc analysis corrected by Sidak’s multiple comparison test indicated that AB-free kava reduced normalized airway elastance at the 100 mg/mL dose of methacholine compared to the Air and TS groups (Fig. 6D). These results suggested TS alone did not influence airway elastance relative to the Air group, but that the combination of TS + AB-free kava dampened airway elastance responses.

We next examined three additional airway properties in response to broadband frequency forced oscillation: Newtonian resistance, tissue damping, and tissue elastance. Broadband frequency allows for information about airway and tissue responses to a test signal that is above and below the subject’s breathing frequency. When assessing Newtonian resistance, which is the resistance of the central and conducting airways, we found no effect of treatment on basal (Fig. 6E) or methacholine-induced responses (Fig. 6F). We also found no effect of treatment on tissue damping (Fig. 6G and 6H) or on tissue elastance (Fig. 6I, and 6J).

We observed no impact of TS on basal properties or methacholine-induced properties of airway mechanics. The lack of an effect of TS on airway mechanics suggests that exposure to TS was not robust enough or chronic enough to elicit detectable deficits. Interestingly, we did observe that AB-free kava treatment reduced responses to methacholine in two of the airway properties measured: airway elastance and airway resistance. This finding suggests that AB-free kava reduced bronchoconstriction and/or mucus secretion caused by methacholine administration. Though the mechanisms responsible for this effect are unclear, prior studies have shown that β-adrenergic receptor agonists can increase mucus secretion by increasing the number of mucus producing cells.^*79, 80*^ Thus, it is possible that AB-free kava reduced activation of the β-adrenergic receptor or its downstream signaling partners, contributing to its lung function modulation.

### The PKA/CREB/LKB1/COX-2 pathway

Results of our previous studies showed that tobacco specific toxicant, NNK, and its major metabolite, NNAL, can function as agonists of β-adrenergic receptor (AR) to activate the protein kinase A (PKA) signaling pathway, leading to cAMP response element-binding protein (CREB) phosphorylation/activation, liver kinase B1 (LKB1) phosphorylation/deactivation, and COX-2 up-regulations in cells and in mice.^*81, 82*^ CREB phosphorylation has also been reported to be higher in the buffy coats from smokers than non-smokers and such an elevation was associated with smoking addiction behaviors.^*83*^ In addition, CREB activation could result in stress and anxiety^*84*^ while its deletion promotes mental resilience.^*85*^ CREB, as a transcription factor, has also been reported to regulate a panel of downstream genes, including COX-2, which may contribute to inflammation. LKB1 is a tumor suppressor protein and plays a key role in cancer immunology.^*86*^ Its deactivation typically leads to the inhibition of the AMPK pathway and the activation of mTOR. Metformin, a classical AMPK activator, has been recently demonstrated to reduce withdrawal signs precipitated by nicotine.^*87*^ TS exposure thus may modulate this signaling pathway, which may contribute to its various deleterious effects, including those evaluated above. We thus characterized several representative signaling events in this pathway using the lung tissues from Round 2 via Western Blotting analysis (Fig. 7). The 4-week tobacco smoke exposure increased the phosphorylation of LKB1, CREB, and mTOR and reduced the phosphorylation of AMPK, indicating the activation of CREB, mTOR and the deactivation of LKB1 and AMPK. The 4-week tobacco smoke exposure also increased the level of COX-2, which may contribute to the observed lung inflammation. AB-free kava dietary supplementation alone without tobacco smoke exposure had no obvious effects on any of these molecular changes. As expected, AB-free kava dietary supplementation in general neutralized the changes caused by tobacco smoke exposure, reducing tobacco smoke-induced phosphorylation of CREB, mTOR, and LKB1 while restoring AMPK phosphorylation compromised by tobacco smoke exposure. AB-free kava also reduced COX-2 elevation caused by tobacco smoke exposure. These data overall suggest that tobacco smoke exposure may activate the PKA/CREB/COX-2 signaling pathway, potentially responsible for nicotine addiction and lung inflammation while AB-free kava corrects such changes.

**Figure 7.**
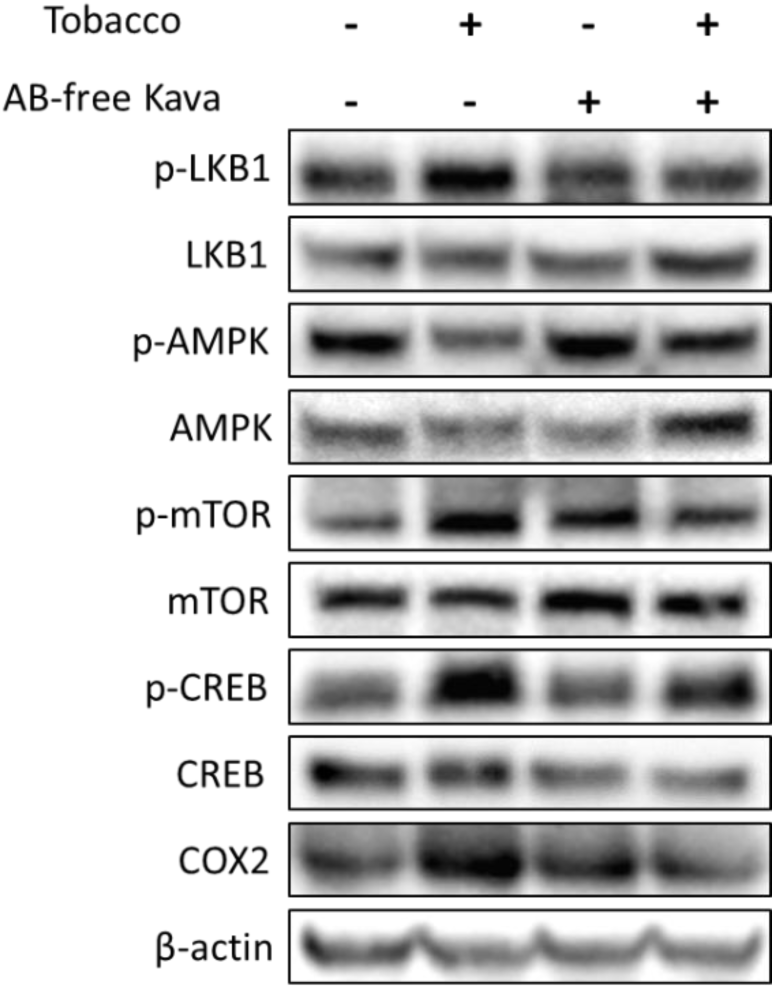
Western blotting analysis of the lung tissues upon TS exposure with/without AB-free kava supplementation, centering on the β-AR mediated PKA/CREB/LKB1/COX-2 pathway.

## CONCLUSIONS

Tobacco smoke (TS) exposure is well accepted as a major serious and long-lasting public health issue, which contributes to the cause of a wide range of health problems, including numerous chronic diseases. Due to the addictive nature of nicotine, an ideal intervention should not only reduce TS dependence/use but also protect individuals from damages caused by TS exposure, particularly those exposed to second hand smoke, to achieve harm reduction. The current study assessed the effects of dietary AB-free kava supplementation on a panel of representative biological functions perturbed by TS exposure in mice. The results from the RNA-Seq analyses of the lung and liver tissues demonstrate that AB-free kava effectively and globally neutralizes the transcriptional perturbation induced by TS exposure in mice. These results, for the first time, reveal the unique potential of AB-free kava to holistically counteract the deleterious effects induced by TS exposure, a result which has yet to be demonstrated by any other known treatments. To substantiate this novel observation, lung inflammation, nicotine withdrawal, lung function, and a potential underlying mechanism – the PKA signaling pathway, were characterized. TS exposure, as expected, induced lung inflammation, resulted in nicotine withdrawal, compromised lung function, and induced the activation of the PKA pathway. Consistent with the RNA-Seq results, AB-Free kava dietary supplementation safely and effective offset these diverse perturbations induced by TS exposure, namely suppressing lung inflammation, ameliorating nicotine withdrawal symptoms, restoring mouse lung function and suppressing PKA activation under the current experimental conditions, which mimics the level of TS exposure among smokers while the daily dosage of AB-free kava is comparable to those recommended for human use.

In conclusion, this is the first demonstration that AB-free kava has the potential to deliver a holistic protective effect against TS exposure in a pre-clinical animal model with high physiological relevance. These results overall are in alignment with the preliminary observations from our earlier pilot smoker clinical trials.^*57*^ The potential impact of this study on improving human health could be tremendous if implemented properly. Future more systematic characterizations of AB-free kava against TS exposure is warranted in additional pre-clinical animal models to characterize the scope, determine the responsible ingredient(s), identify the molecular target(s), elucidate the underlying mechanisms, and fill in other gaps before its clinical translation.

## ASSOCIATED CONTENT

### Supporting Information

Fig. S1. HPLC traces of AB-free kava and that recovered from the diet with the estimated abundance of the six major kavalactones based on the absorbance at 240 nm.

Fig. S2. Bodyweight changes of mice in Round 1 during the 4-week experimental period.

Table S1. Information of antibodies used in Western Blotting analyses.

## AUTHOR INFORMATION

### Author Contributions

**Tengfei Bian** – Department of Medicinal Chemistry, Center for Natural Products, Drug Discovery and Development (CNPD3), College of Pharmacy, University of Florida, Gainesville, Florida 32610, United States

**Allison Lynch** – Department of Medicinal Chemistry, Center for Natural Products, Drug Discovery and Development (CNPD3), College of Pharmacy, University of Florida, Gainesville, Florida 32610, United States

**Kayleigh Ballas** – Department of Medicinal Chemistry, Center for Natural Products, Drug Discovery and Development (CNPD3), College of Pharmacy, University of Florida, Gainesville, Florida 32610, United States

**Jessica Mamallapalli** – Department of Medicinal Chemistry, Center for Natural Products, Drug Discovery and Development (CNPD3), College of Pharmacy, University of Florida, Gainesville, Florida 32610, United States

**Breanne Freeman** – Department of Medicinal Chemistry, Center for Natural Products, Drug Discovery and Development (CNPD3), College of Pharmacy, University of Florida, Gainesville, Florida 32610, United States

**Alexander Scala** – Department of Medicinal Chemistry, Center for Natural Products, Drug Discovery and Development (CNPD3), College of Pharmacy, University of Florida, Gainesville, Florida 32610, United States

**Yifan Wang** – Department of Medicinal Chemistry, Center for Natural Products, Drug Discovery and Development (CNPD3), College of Pharmacy, University of Florida, Gainesville, Florida 32610, United States

**Hussein Trabouls** – Division of Experimental Medicine McGill University Health Center, 1001 Decarie Boulevard, Montreal, Qc, H4A3J1

**Ranjithkumar Chellian** - Department of Psychiatry, College of Medicine, University of Florida, Gainesville, Florida 32610, United States

**Amy Fagan** - Department of Physiological Sciences, College of Veterinary Medicine, University of Florida, Gainesville, Florida 32610, United States

**Zhixin Tang** - Department of Biostatistics, College of Public Health and Health Professionals & College of Medicine, University of Florida, Gainesville, Florida 32610, United States

**Haocheng Ding** - Department of Biostatistics, College of Public Health and Health Professionals & College of Medicine, University of Florida, Gainesville, Florida 32610, United States

**Umasankar De** - Department of Pathology, Immunology and Laboratory Medicine, College of Medicine, University of Florida, Gainesville, Florida 32610, United States

**Kristianna M. Fredenburg** - Department of Pathology, Immunology and Laboratory Medicine, College of Medicine, University of Florida, Gainesville, Florida 32610, United States

**Zhiguang Huo** - Department of Biostatistics, College of Public Health and Health Professionals & College of Medicine, University of Florida, Gainesville, Florida 32610, United States

**Carolyn J. Baglole** - Division of Experimental Medicine McGill University Health Center, 1001 Decarie Boulevard, Montreal, Qc, H4A3J1

**Weizhou Zhang** - Department of Pathology, Immunology and Laboratory Medicine, College of Medicine, University of Florida, Gainesville, Florida 32610, United States

**Leah R. Reznikov** - Department of Physiological Sciences, College of Veterinary Medicine, University of Florida, Gainesville, Florida 32610, United States

**Adriaan W. Bruijnzeel** - Department of Psychiatry, College of Medicine, University of Florida, Gainesville, Florida 32610, United States

**Chengguo Xing** - Department of Medicinal Chemistry, Center for Natural Products, Drug Discovery and Development (CNPD3), College of Pharmacy, University of Florida, Gainesville, Florida 32610, United States

### Notes

The manuscript was written through contributions of all authors. All authors have given approval to the final version of the manuscript.

### Notes

Any additional relevant notes should be placed here.

## ACKNOWLEDGMENT

We thank the University of Florida Department of Medicinal Chemistry for its support with LC/MS analyses.

## ABBREVIATIONS

TS: Tobacco smoke
TSP: total suspended particles
BAL: bronchoalveolar lavage
PKA: Protein Kinase A
CREB: cAMP Response Element-Binding Protein
LKB1: Liver Kinase B1
PCA: Principal Component Analysis
IPA: Ingenuity Pathway Analysis

