## Supplemental data for "AB-free kava enhances resilience against the adverse health effects of tobacco smoke in mice"

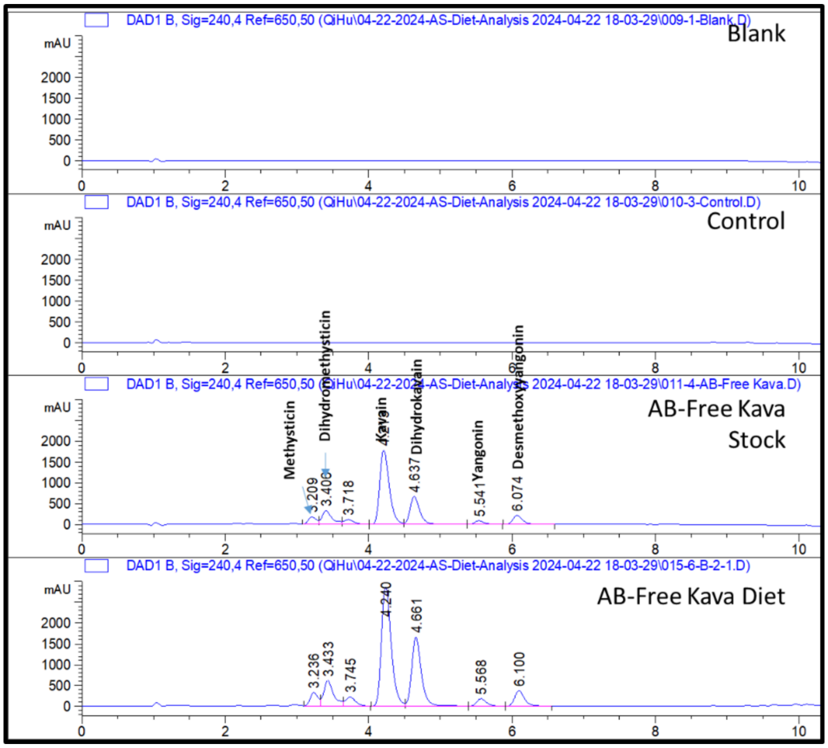


Fig. S1. HPLC traces of the original AB-free kava stock and AB-free kava recovered from the diet with associated background traces.

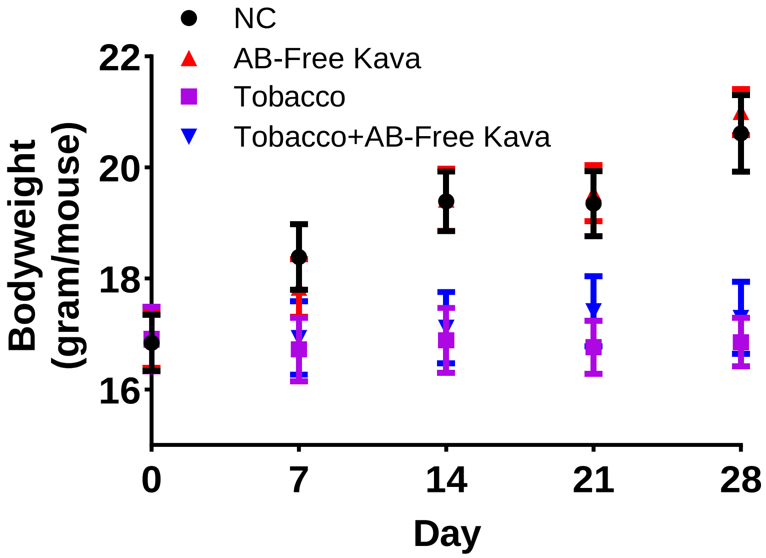


Fig. S2. Bodyweight changes during the 4-week experimental period among the four groups of mice (Mean ± SE).

Table S1. Information of Antibodies used for Western Blotting analyses in this study

| Antibodies | Company | Catalog number |
| --- | --- | --- |
| LKB1 (D60C5) Rabbit mAb | Cell Signaling Technology | 3047 |
| Phospho-LKB1 (Ser428) (C67A3) Rabbit mAb | Cell Signaling Technology | 3482 |
| CREB (48H2) Rabbit mAb | Cell Signaling Technology | 9197 |
| Phospho-CREB (Ser133) (87G3) Rabbit mAb | Cell Signaling Technology | 9198 |
| AMPKα Antibody | Cell Signaling Technology | 2532 |
| Phospho-AMPKα (Thr172) (40H9) Rabbit mAb | Cell Signaling Technology | 2535 |
| mTOR (7C10) Rabbit mAb | Cell Signaling Technology | 2983 |
| Phospho-mTOR (Ser2448) (D9C2) XP® Rabbit mAb | Cell Signaling Technology | 5536 |
| Cox2 (D5H5) XP® Rabbit mAb | Cell Signaling Technology | 12282 |
| Anti-rabbit IgG, HRP-linked Antibody | Cell Signaling Technology | 7074 |
| Mouse monoclonal anti-beta-actin | Sigma Aldrich | A2228 |
